# Asymmetric perirenal brown adipose dormancy in adult humans is defined by local sympathetic activity

**DOI:** 10.1101/368621

**Authors:** Naja Z. Jespersen, Amir Feizi, Eline S. Andersen, Sarah Heywood, Helle B. Hattel, Søren Daugaard, Per Bagi, Bo Feldt-Rasmussen, Heidi S. Schultz, Ninna S. Hansen, Rikke Krogh-Madsen, Bente K. Pedersen, Natasa Petrovic, Søren Nielsen, Camilla Scheele

## Abstract

We here detect dormant brown adipose tissue (BAT) in adult humans, occurring in most of the perirenal fat depot and characterized by a unilocular morphology. This phenotype was contrasted by multilocular BAT accumulating near the adrenal gland. Transcriptomic analysis revealed a gene expression profile of unilocular BAT that was approaching, yet was still distinct from, the expression profile of subcutaneous white adipose tissue (WAT). Candidate gene signatures were recapitulated in a murine model of unilocular brown fat induced by thermoneutrality and high fat diet. We identified SPARC as a candidate adipokine representing a dormant BAT state in the absence of sympathetic activation and CLSTN3 as a novel marker for multilocular BAT. Brown fat precursor cells were present in the entire perirenal fat depot, regardless of state. When differentiated in vitro, these cells responded to acute norepinephrine stimulation by increasing UCP1 gene expression and uncoupled respiration, confirming a BAT phenotype. We thus propose a mechanism for the reduction of functionally competent BAT in adult humans and we provide a solid data set for future research on factors that can reactivate dormant BAT as a potential strategy for combatting obesity and metabolic disease.

## Introduction

Metabolically active brown adipose tissue (BAT) was discovered a decade ago in adult humans in the supraclavicular and deep neck region, along the spinal cord and around the kidneys (the perirenal adipose depot), using PET/CT-scan technology^1–6^. BAT provides non-shivering thermogenesis through mitochondrial uncoupling via the BAT specific protein, uncoupling protein 1 (UCP1)^7^. This is an energy consuming process, thus providing a rationale for targeting human BAT as part of an anti-obesity strategy. In accordance, the amount of active human BAT has been found to negatively correlate with body mass index (BMI)^3^ and to increase metabolic rate^8^. An additional beneficial property of BAT in promoting metabolic fitness, is its glucose uptake capacity, and several studies have demonstrated an increase in insulin sensitivity upon increased BAT activity in adult humans^9–12^.

In larger studies of adult humans, the frequency of cold-activated BAT (as detected with PET/CT-scans) has been described to be around 44% ^3, 13–15^. However, active BAT is less frequently found in older compared to younger individuals, and only approximately 14% of the subjects over 40 years of age appear to have BAT that can be activated by cold^3,13,14^. It can therefore be questioned whether BAT activation would be an efficient approach to treat metabolic dysfunction in middle-aged and elderly humans. On the other hand, current methods for in vivo BAT quantifications are limited to detection of activated BAT, and the actual amount of the tissue might therefore be underestimated.

An age-dependent shift in BAT morphology was described in early autopsy-histology studies of human BAT, demonstrating that a more white-like phenotype gradually occurred with age, represented by increased lipid accumulation^16,17^. It has been suggested that adult human BAT is more comparable to the inducible type of BAT, the so-called beige or brite fat^18–21^. This idea is based on its heterogeneity in morphology as well as in marker gene expression in comparison to murine classical brown and beige fat depots^22,23^. However, it is currently elusive whether the infant BAT depots are gradually replaced with white fat or if it retains a BAT identity while acquiring a more white-like fat phenotype. The latter kind of fat state might be more efficiently convertible into active BAT compared to classical white fat tissue.

Whereas the supraclavicular depot is the most characterized BAT depot in human adults, UCP1 expression has also been detected in the perirenal fat depot in samples collected near the hilus and adjacent to retroperitoneal and adrenal gland tumors.^24–29^. Perirenal fat surrounds the kidney and the adrenal gland, which both have extensive, but separate blood supplies. The autonomic innervation of the kidney is entirely through the sympathetic nervous system (SNS), with nerve bundles docking at the hilus. The adrenal gland can be considered an extension of the SNS and it produces and secretes epinephrine and norepinephrine (NE) in response to sympathetic stimulation^30^. Patients suffering from NE-producing adrenal gland tumors, so called pheochromocytomas, can exhibit massive browning of the adjacent visceral fat ^27,31,32^. This is most pronounced in perirenal fat and strongly suggests that NE originating from the adrenal gland can induce adipose browning^33–35^. In the current study we therefore hypothesized that a multilocular BAT phenotype would exist close to local sources of sympathetic activation, i.e. the adrenal gland. We also sampled at distant locations within the perirenal fat depot, with the aim of investigating the occurrence of dormant BAT. Therefore, we characterized the BAT phenotype of perirenal adipose biopsies from four different regions surrounding the kidney and adrenal gland and included subcutaneous fat from the incision site as a control.

## Results

We recruited 20 kidney donors, aged 37-68 years, to investigate the status of perirenal fat, a depot consisting of BAT during infancy. The subjects were healthy and included both men (n=11) and women (n=9). The physiological characteristics of the subjects are summarized in Table 1.

**Table 1.** Subject characteristics.

| **Sex**<br>**(N)** | **Age**<br>**(years)** | **BMI**<br>**(kg/m<sup>2</sup>)** |
| --- | --- | --- |
| **Men**<br>**(11)** | 55<br>(38 – 68) | 26<br>(24 – 37) |
| **Women**<br>**(9)** | 54<br>(37 – 67) | 25<br>(16 – 28) |

Data are median and range. Abbreviations: BMI; Body mass index.

### The β3-adrenergic receptor and BAT markers characterizing perirenal BAT

To characterize potential regional differences within the perirenal fat and in relation to white fat (WAT), we investigated the BAT gene expression signature of four perirenal regions including the upper kidney pole, the hilus, the convexity, and the lower kidney pole in a cohort of healthy adults (median age 55 years) (Figure 1A). As a reference tissue, we included subcutaneous fat from the surgical incision site at the abdomen (n=20). As the β3-adrenergic receptor (ADRB3), is responsible for the main BAT activation pathway in rodents^7^, we addressed whether there were any differences in adrenergic receptor gene expression between the perirenal regions. We performed qPCR mapping of the gene expression of nine different α- and β-adrenergic receptors in the four perirenal regions. We included supraclavicular BAT with high UCP1 expression (n=6) ^22^ and subcutaneous WAT for comparison. We found that the β3-adrenergic receptor (ADRB3), was substantially higher in *all* perirenal regions compared to the subcutaneous adipose tissue but was not different from the supraclavicular BAT (Figure 1B). Furthermore, the β1-adrenergic receptor (ADRB1) was higher in some perirenal regions and in supraclavicular fat, while the β2-adrenergic receptor (ADRB2) was higher in subcutaneous fat compared to all other regions (Figure 1B). Both ADRB3 and ADRB1 mRNA expression levels were highly correlated to UCP1 in a multivariate linear regression analyses (P < 0.0001, β-coefficients = 0.39 and 1.33 respectively), while there was a strong negative correlation with ADRB2 (P < 0.0001, β-coefficient = - 0.89). This is interesting, given that the β1-adrenergic receptor has been demonstrated to perform BAT activation in the absence of β3-adrenergic receptors^36,37^ while the role of the β2-adrenergic receptor remains to be explored. The model was overall significant (P < 0.0001 and adjusted R^2^ = 0.423). None of the α-adrenergic receptors were significantly correlated to UCP1 mRNA expression. Thus, based on adrenergic receptor gene expression data, all perirenal regions seemed to have equal expression of the main receptor for BAT activation; which was comparable with supraclavicular BAT and distinct from subcutaneous WAT.

**Figure 1.**
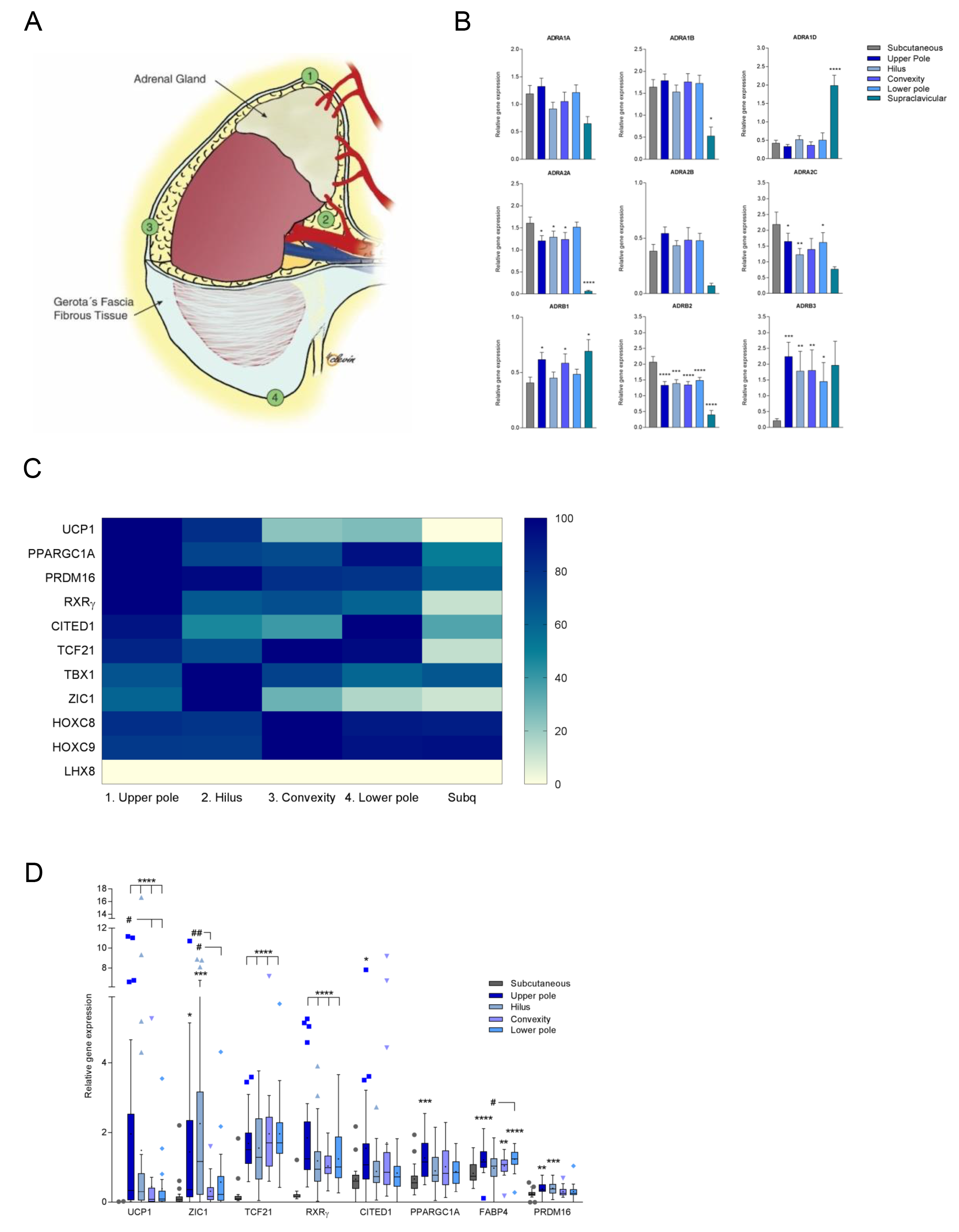
qPCR profiling of adrenergic receptors and brown fat markers in perirenal adipose. **(A)** An illustration of the human kidney. Numbers label the four perirenal regions from where surgical biopsies were obtained; 1= the upper kidney pole, 2 = the hilus, 3 = the convexity and 4 = the lower kidney pole. A subcutaneous fat biopsy was obtained from the incision site at the abdomen. **(B)** Distribution of adrenergic receptors (AR) in human perirenal, supraclavicular and subcutaneous fat. Data are presented as means (bars) with whiskers representing standard error of the mean (SEM). Comparisons were calculated between subcutaneous fat and perirenal / supraclavicular fat using a mixed model ANOVA and subcutaneous fat as a control region, except for the comparison between ADRA1A, ADRA1D, ADRA2A, ADRA2B and ADRA2C in supraclavicular and subcutaneous fat. Due to infinite likelihood, a mixed model ANOVA could not be applied for this comparison and thus a one-way ANOVA was performed to assess differences between supraclavicular and subcutaneous expression of these receptors. A P-value below 0.05 was considered statistically significant. * = significant difference between subcutaneous fat and perirenal / supraclavicular regions. * = P < 0.05, **= P < 0.01, ***= P < 0.001, **** = P < 0.0001. Subcutaneous N = 20, upper pole = 29, hilus = 30, convexity = 19, lower pole = 23 and supraclavicular = 6. **(C)** Heatmap illustrating relative expression levels of all investigated genes. Regions are sorted based on UCP1 expression. **(D)** Perirenal selective genes, i.e. genes that display higher mRNA levels in fat from one or more perirenal regions compared to subcutaneous fat. Data are presented as Tukey box plots with boxes representing the interquartile range (IQR), whiskers = 25 - 75 % percentile and • = Mean. Values > 1.5 times x IQR are marked as individual data points. Comparisons were calculated using mixed models ANOVA with Tukey correction. * = significantly different from subcutaneous fat. # = significant difference between perirenal regions. * = P < 0.05, **= P < 0.01, ***= P < 0.001, **** = P < 0.0001. See also Figure S1. Subcutaneous N = 20, upper pole = 29, hilus = 30, convexity = 19 and lower pole = 23.

We next investigated the gene expression of brown, brite/beige-, white-fat markers in the four perirenal regions specified in Figure 1A, and subcutaneous fat from the incision site. The selected markers were previously identified in murine studies and used for characterization of human supraclavicular fat ^20–22, 38^. For some subjects, multiple samples were collected from each region. In these cases, the sample with the highest level of UCP1 was chosen for analysis of all markers in the initial analysis, thus including the upper kidney pole (n=18), the hilus (n=20), the convexity (n=16), and the lower kidney pole (n=17). We found the highest UCP1 expression in the upper pole (the region closest to the adrenal gland), followed by the hilus (the main site of sympathetic innervation), the convexity, and finally the lower pole (Figure 1C). Thus, the two regions closest to NE trafficking had a more pronounced BAT signature.

We next compared the expression levels of the markers between the four perirenal regions and the subcutaneous fat. Here, we included all samples, thus multiple samples for most subjects. In total, 121 tissue samples were included in the analysis. UCP1 was expressed to a higher degree in all perirenal regions compared to subcutaneous fat (Figure 1D) and UCP1 mRNA levels at the upper kidney pole were higher than at the lower pole and the convexity (Figure 1D). Interestingly, also the recently established human perirenal BAT marker RXRγ^28^ displayed increased expression in all the perirenal regions compared to subcutaneous fat (Figure 1D). In accordance with previous studies, UCP1 was undetectable or barely detectable in the subcutaneous samples^21,22^ (Figure 1D and Table S1). The BAT activity genes, PPARGC1A^39^ and CITED1^22,38^, were higher expressed in the perirenal upper pole only when comparing the perirenal regions to subcutaneous fat (Figure 1D). ZIC1 has been suggested to be a prominent marker of the so-called classical BAT in rodents^40^ and has frequently been measured for characterization of human BAT, although the expression is low in the supraclavicular region of adult humans^21,22^. In the current study, ZIC1 as well as PRDM16 were expressed at higher levels both by the upper pole and at the hilus when comparing to subcutaneous fat, and ZIC1 expression was higher in the hilus area than in the region of the convexity and the lower pole when comparing perirenal fat regions (Figure 1D). Whereas human perirenal and supraclavicular BAT have several markers in common, some markers previously described as human BAT selective (LHX8 and TBX1) or white fat selective (HOXC8 and HOXC9) (Figure 1D and Figure S2) ^20,22^, were not differentially regulated between subcutaneous and perirenal fat. In addition, TCF21, a murine visceral white fat marker ^41^ was higher expressed in the perirenal regions than in the subcutaneous depot (Figure 1D). Thus, our data indicate that the molecular signature of perirenal fat differs from that of supraclavicular and that perirenal BAT is sympathetically regulated with a higher proportion of active BAT in the regions close to the adrenal gland.

### UCP1-positive unilocular BAT dominates perirenal fat in adult humans

We next assessed the morphology and UCP1 protein distribution in the four perirenal regions specified in Figure 1A. Immunohistochemistry performed on perirenal fat revealed two distinct types of UCP1 positive adipocytes; one with the morphological resemblance of classical brown adipocytes, i.e. multilocular BAT (MBAT) and one with a unilocular BAT morphology (UBAT+) (Figure 2A). A third type of adipocytes which were unilocular and UCP1 negative were also detected and classified as UBAT-(Figure 2A). Importantly, unilocular UCP1 positive adipocytes have previously been observed in human perirenal fat^28^. In total, 20% of all samples were classified as MBAT, and MBAT was detected in 60% of the participants. In most cases MBAT sections also included areas classified as UBAT+. Importantly, UBAT+ was observed in nearly all perirenal samples of the dataset, while only 1.8% of the samples were completely negative for UCP1 (UBAT-) (Figure 2B). Both UBAT+ and UBAT-areas were often found in distinct, but adjacent islets (Figure S2A). A positive and negative control for the staining was included (Figure S2B). Interestingly, MBAT positive samples were observed at highest frequency by the upper kidney pole (30%), followed by the hilus (25%), the convexity (15.7%) and the lower pole (3.6%) (Figure 2B). UBAT+ was in contrast found in all subjects and all perirenal regions. To further address this finding, we stained for two additional BAT markers; PRDM16 and RXRγ. PRDM16 was previously found to be associated with UCP1 expression^42–44^ and was selective for brown fat progenitors isolated from the supraclavicular depot of adult humans^22^, while RXRγ previously was described as a perirenal fat marker which clustered with UCP1 expression^28^. Immunohistochemical stainings of RXRγ were positive in all perirenal samples (Figure S2B upper panel), whereas 88 % of perirenal fat samples were positive for PRDM16 (Figure S2B lower panel). No selection towards MBAT positive areas was observed for either RXRγ or PRDM16 positivity. To further investigate the UCP1 unilocular adipocytes, we performed western blot analyses on a selection of tissue samples derived from biopsies where the corresponding histology was either multilocular or unilocular. Although the adjacent piece of the biopsy used for western blot analysis could differ from the piece used for histology, we hereafter refer to these samples as “multilocular” or “unilocular” based on the histology data. We also included subcutaneous samples to investigate whether the unilocular samples would be more similar to white fat than to multilocular fat. Despite variation between individual samples, we could confirm that unilocular samples were indeed positive for UCP1 protein (Figure 2C, left panel). To obtain an indication on the association between sympathetic activation and the multilocular phenotype, we measured tyrosine hydroxylase which was higher in the multilocular samples, suggesting higher sympathetic activation (Figure 2C, right panel), in line with previous morphological observations^32^. To assess potential differences in mitochondrial activity, we measured a panel of OXPHOS proteins and observed increased expression of mitochondrial complexes III and IV in perirenal compared to subcutaneous fat. However, similarly to the UCP1 protein expression, there was no difference in OXPHOS protein levels between unilocular and multilocular samples (Figure 2D).

**Figure 2.**
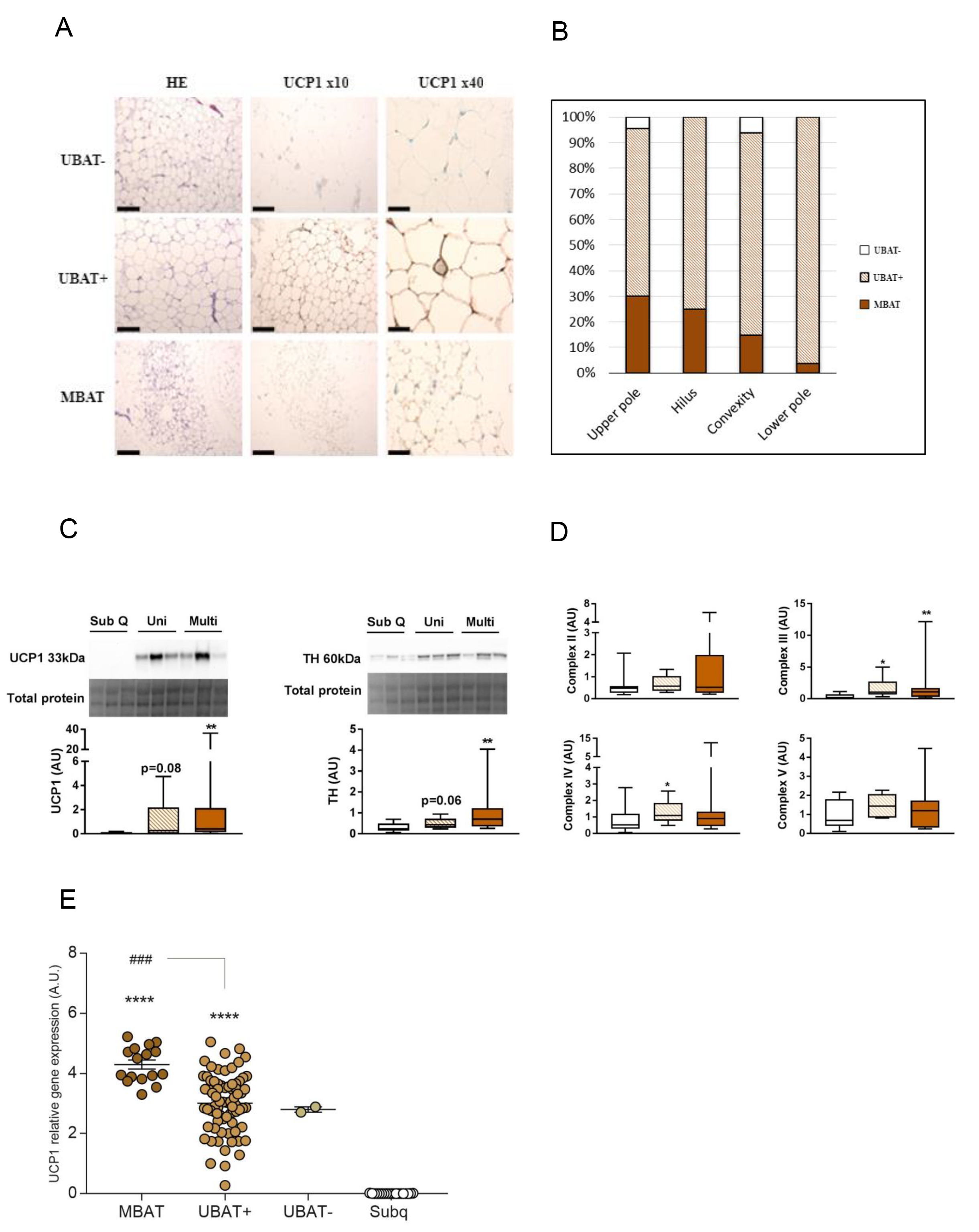
The brown fat phenotypes in perirenal tissue of adult humans. **(A)** Morphological and immunohistochemical evaluation of human perirenal fat. Representative H&E staining (left panel) and UCP1 staining (central and right panel) of areas with multilocular UCP1 positive brown adipose tissue (MBAT) (lower row) unilocular BAT (UBAT+) (middle row),, and unilocular UCP1 negative BAT (UBAT-) (upper row).;. Bar scale = 240 μm in left and middle panel and 60 μm in right panel. **(B)** Percentage distribution of MBAT, UBAT+ and UBAT-samples in the different perirenal regions. **(C)** Multilocular and unilocular perirenal fat and subcutaneous fat samples were analyzed for UCP1 and Tyrosine hydroxylase (TH) protein expression using western blot. A representative band from the total protein assessment is shown and was used for quantification. **(D)** Multilocular and unilocular perirenal fat and subcutaneous fat samples were analyzed for the mitochondrial OXPHOS complexes. Complex I was too weak to quantify. All blots are shown in the supplementary information (Fig. S2D). **(E)** UCP1 mRNA expression in groups made from histology data, i.e. divided into MBAT, UBAT+ and UBAT- and with the corresponding subcutaneous samples as reference material (Subq). Data are mean +/-SE; *P < 0.05, **P < 0.01, ***P < 0.001.

We now went back to the UCP1 mRNA data described in Figure 1 and divided our sample set into multilocular and unilocular based on the histology data. We observed that a high UCP1 mRNA expression overall seemed to coincide with a multilocular morphology (Figure 2E). We next sought to address how the other adipose type markers were related to UCP1 expression in the two morphological groups. We thus performed multivariate regression analyses including all samples and all investigated markers while applying manual backwards elimination of independent variables (Table 2). In concordance with the gene signature in supraclavicular fat^22^, this analysis demonstrated that in unilocular samples, UCP1 correlated positively with the master regulator of mitochondrial biogenesis, PPARGC1A (P < 0.05). Moreover, we observed a negative correlation with HOXC8 (P < 0.05) a gene described as a white fat marker^22,41,45^. In addition, we confirmed previous reports identifying RXRγ as a perirenal BAT marker^28^ demonstrating a positive correlation with UCP1 (P < 0.0001). The same pattern was observed in the multilocular samples where UCP1 correlated negatively with HOXC8 (P < 0.05) and positively with RXRγ (P < 0.05). None of the other measured markers correlated to UCP1 in neither uni-nor multilocular samples (Table 2). Taken together, these results raised the possibility of a large pool of dormant BAT, distinct from subcutaneous fat, being present in the entire perirenal fat depot, even in subjects around 60 years of age.

**Table 2.** Genes correlated to UCP1 in unilocular and multilocular tissue samples.

| **Genes** | **morphology** | **P-value** | **β-coefficient** |
| --- | --- | --- | --- |
| **UCP1 vs. RXRγ** | Uni | < 0.0001 | 1.52 |
| **UCP1 vs. PPARGC1A** | Uni | < 0.05 | 0.36 |
| **UCP1 vs. HOXC8** | Uni | < 0.05 | -0.47 |
| **UCP1 vs. HOXC8** | multi | < 0.05 | - 1.37 |
| **UCP1 vs. RXRγ** | multi | < 0.05 | 1.29 |

### The gene signature of human dormant BAT

Based on our results, we concluded that the regions presenting unilocular adipocytes, which were positive for UCP1, represented a state of dormant BAT. To validate this conclusion and further describe the molecular signature of this state, we performed RNA sequencing on a subset of multilocular, unilocular and subcutaneous samples (n=4 for each adipose tissue type). The principal component analysis (PCA) of the transcriptome readouts, including all identified genes in all samples, demonstrated a separation into three subpopulations. These subpopulations were, with the exception of one subject, defined by the origin of the samples (i.e. subcutaneous, unilocular or multilocular) (Figure 3A). Next, we performed differentially expression analysis between the samples using the grouping information from PCA results. Here, we used neighbour joining algorithms to cluster samples based on statistically significant differentially expressed genes (FDR < 0.01). This resulted in fat tissue type-dependent clustering of the multilocular, unilocular and subcutaneous groups, suggesting that these groups were defined by unique gene expression signatures (Figure 3B). We observed both up- and down-regulated genes for all three comparisons; unilocular versus multilocular, subcutaneous versus unilocular and subcutaneous versus multilocular (Figure S3A)). Strikingly, several mitochondrial genes were higher expressed in multilocular compared to both unilocular and subcutaneous, these included the mitochondrially transcribed: MT-CO1, MT-CYB, MT-ATP6, MT-ND4, MT-ND1, MT-CO2, MT-CO3. Moreover, genes transcribed from the nucleus with mitochondria-associated gene products were higher, including the brown fat specific UCP1 and its transcriptional coactivator PPARGC1A. In concordance with previous studies^46,47^, various forms of a key protein in mitochondrial respiration, Creatine kinase, was also higher in the multilocular samples. Taken together, these observations suggest a higher metabolic activity in the multilocular fat compared to the subcutaneous and the unilocular fat (Figure 3B). To explore the differentially expressed genes between multilocular and unilocular in higher resolution, we used a volcano plot (based on −log_10_ p-values versus log_2_ fold change). This revealed CLSTN3 as the most significantly regulated gene between the two conditions, upregulated in biopsies with a multilocular BAT morphology (Figure 3C). Other top candidates accumulating in this group included TBATA (encoding Protein TBATA), CYP1A2 (encoding Cytochrome P450 1A2), ZDHHC19 (encoding Probable palmitoyltransferase ZDHHC19) and KCNK3 (encoding Potassium channel subfamily K member 3). Interestingly, KCNK3 was previously reported to be higher expressed in immortalized human brown adipocyte compared to white adipocytes, and to be important for thermogenic function^48^, supporting our idea that the multilocular adipose samples represented a more active BAT phenotype compared to the unilocular adipose samples.

**Figure 3.**
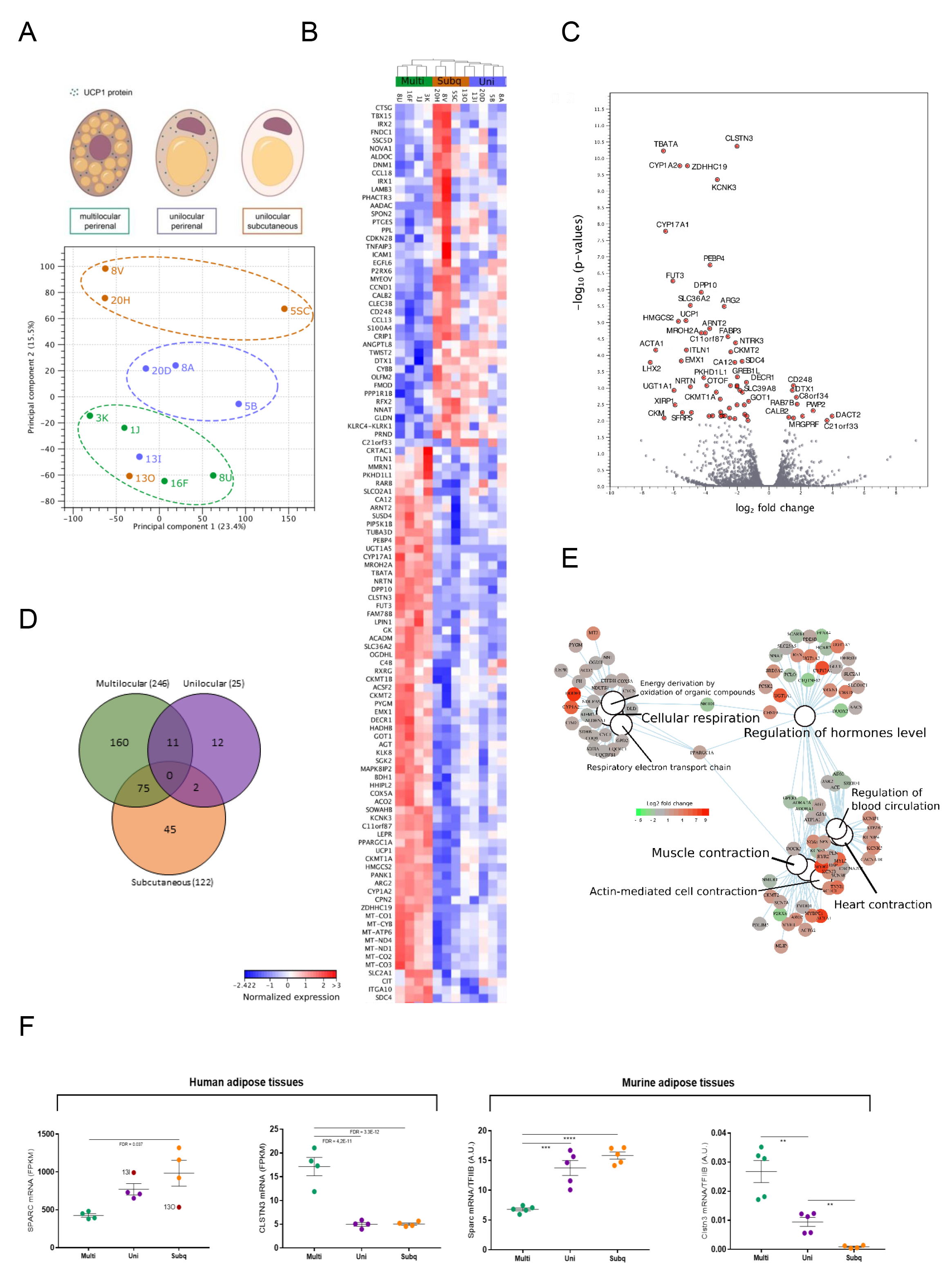
RNA sequencing of multilocular and unilocular perirenal fat and subcutaneous fat. **(A)** Principal component analysis (PCA) plot demonstrating the separation based on transcriptome profiles of the three sample-sets including multilocular perirenal, unilocular perirenal and unilocular subcutaneous adipose tissue, with a clear outlier represented by subject 13. **(B)** Heatmap showing clustering of the most differentially expressed genes (FDR < 0.01) between the three tissue types. The expression values are normalized using TMM method (trimmed mean of M values)^69^ to calculate the effective libraries sizes followed by z-score calculation. The color code of the normalized expression values is located next to the heatmap. **(C)** Volcano plot ranking genes after - log10 p-value (y-axis) and log2 fold change (x-axis). The significant genes (p-value < 0.01) are shown with red dots and labeled with their corresponding gene names. **(D)** Venn diagram demonstrating the number of specific/overlaps of differentially expressed genes (FDR < 0.01) for each of the three tissue types. **(E)** Category netplot generated by the clusterProfiler R package^66^. This network plot shows the relationships between the genes associated with the top most significant GO terms (q-value < 0.05) and their corresponding significant fold changes (FDR < 0.01) from multilocular perirenal vs. unilocular perirenal comparison. The log2 foldchange color code is next to the network and the size of the GO terms reflects the q-values of the terms, with the more significant terms being larger. **(F)** *Left panel:* Regulation of candidate genes differentially expressed between groups: multilocular perirenal fat (multi); unilocular perirenal fat (uni) and subcutaneous fat (subq), identified from volcano plot (CLSTN3) or from dispersion ranking between multilocular and subcutaneous to account for gene expression levels (SPARC). Values from the outlier subject 13 (annotated in the graph) are not included in the statistical analysis. Data are adjusted gene counts from the RNA sequencing analysis (FPKM) and adjusted p-value (FDR) is annotated in the figure. *Right panel:* Interscapular BAT (IBAT) was dissected from mice housed at room temperature on chow diet, resulting in multilocular BAT (multi) and from mice housed at thermoneutrality on high fat diet resulting in unilocular BAT (uni). Inguinal WAT was also harvested from the mice housed at thermoneutrality on high fat diet resulting in unilocular WAT (uni). Gene expression analysis of Sparc and Clstn3 was performed by using qPCR. Data are mean +/- SE; Unpaired t-tests between multi IBAT and uni IBAT: *P < 0.05, **P < 0.01, ***P < 0.001. Paired t-tests between uni IBAT and uni IWAT ^$^P < 0.05, ^$$^P < 0.01, ^$$$^P < 0.001

To summarize the number of the specific and overlapping differentially expressed genes among sample groups, we used a Venn diagram with log_2_ fold change cut off of 2 and FDR <0.01 (Figure 3D). As a result, among the differentially expressed genes we noticed that most of them were specific for each analysed tissue (multilocular with the highest number) and we did not find any overlapping genes among the tissues. We next performed gene ontology analysis on the genes differentially expressed between multilocular and unilocular adipose samples (Figure 3E), and between multilocular and subcutaneous adipose samples and between unilocular and subcutaneous adipose samples (Figure S3B). This analysis further emphasized a major difference in cellular respiration by enrichment of the GO terms linked with electron transport chain, actually based on a separate gene-set from what was described above. Interestingly, we further observed a shift in genes regulating hormonal levels. Adipose tissue has an established role in hormonal regulation while our findings raise the idea of differential roles in hormonal regulation of BAT and dormant BAT. Finally, we found a difference in genes involved in muscle contraction. This was surprising to us but might be related to the increased mitochondrial activity combined with the established developmental link between brown adipose and skeletal muscle^41,42^.

To identify additional candidate genes involved in the function or regulation of dormant BAT, we screened for genes with largest fold changes differences between the two most distinct groups, subcutaneous and multilocular. To take the gene expression levels and data dispersion between groups into account, we sorted after standard deviation values. Among the top three genes, we identified SPARC (Adj. p-value: 0.037), a secreted protein and candidate adipokine, which previously was shown to influence and interact with extracellular matrix to regulate cell growth and differentiation^49^. SPARC further caught our interest as we in a parallel study have identified this gene to be higher expressed in WAT compared to supraclavicular BAT (unpublished data). We plotted the RPKM values for SPARC and, consistent with our parallel study, observed a lower expression in multilocular compared to subcutaneous samples. SPARC expression in the unilocular samples accumulated in between multilocular and subcutaneous samples. CLSTN3, the top candidate gene from the Volcano plot, was also plotted, displaying a low expression in both unilocular and subcutaneous compared to multilocular (Figure 3F, left panel). Both marker genes supported a gene signature where the dormant BAT state was approaching the subcutaneous WAT profile.

To further validate the concept of dormant BAT and our candidate genes from adult humans, we utilized a previously established mouse model in which unilocularity was induced in classical BAT by long-term thermoneutrality and high-fat diet^50,51^. We measured the two candidate genes from the human data set in intrascapular BAT from mice kept at room temperature, representing mild cold stimulation for a mouse^51^, and on chow diet (multilocular BAT) and from mice kept at thermoneutrality and on high-fat diet (unilocular BAT) and in inguinal WAT from mice kept at thermoneutrality and on high-fat diet (unilocular WAT)^50,52^. Convincingly, we found that the gene expression pattern of Sparc and Clstn3 in the mouse model was consistent with the pattern observed in the human sample set, supporting the existence of dormant BAT in the perirenal fat of adult humans (Figure 3F, right panel). Interestingly, SPARC has been shown to have an opposing expression pattern to UCP1 in hibernating arctic ground squirrels^53^. This could suggest a suppressive role of SPARC in relation to active BAT. Calsyntenin-3 belongs to a family of post-synaptic membrane proteins shown to be highly expressed in GABAergic neurons^54^. The precise role of this protein in active versus dormant BAT or WAT remains to be investigated. Taking the human and the mouse data together, we identify SPARC as a novel marker for unilocular, i.e. dormant BAT and CLSTN3 as a marker for multilocular BAT, in mammals.

### Brown fat precursor cells are present in all perirenal regions

To further characterize the multilocular and unilocular BAT in the perirenal fat depot, we isolated adipogenic progenitor cells from the different perirenal fat regions and from subcutaneous abdominal fat using our previously established protocol for human brown and white adipocytes^22^. To assess the purity of the cultures we examined their surface marker expression by flow cytometry analysis. When characterizing the cultures from the four regions (n = 4 from each region) we found the cultures to be negative for CD31 (endothelial marker) and CD45 (hematopoietic stem cell marker) while staining positive for CD90 and CD166, demonstrating high adipogenic potential of the cells (Figure 4A)^55^. For CD90, cell cultures were classified as highly positive or intermediately positive. There were more intermediately positive cells in the hilus compared to the subcutaneous derived cultures (Table S1). However, most cells were still highly CD90 positive, and the slightly increased number of intermediately vs. highly CD90 positive cells was not considered to impact the results. In support, no difference in adipogenic potential was observed between the groups (Figure S4). To characterize the cells further, we differentiated adipogenic progenitors isolated from the four perirenal regions as well as from the subcutaneous depot (n=18). On the final day of differentiation, mature lipid droplet containing adipocytes were stimulated with either 10 μM of norepinephrine (NE) or vehicle treatment for four hours and were then harvested for RNA isolation and gene expression analyses. A striking UCP1 up-regulation in response to four hours of NE treatment was observed in cells derived from all perirenal fat regions. This was in contrast to the cells isolated from the subcutaneous depot (Figure 4B). PPARGC1A, which has been shown to be robustly increased in BAT upon cold-induced activation^39,40^ was also induced by NE treatment in all perirenal groups except for the lower kidney pole, while no PPARGC1A induction was observed in the subcutaneous adipocytes (Fig 4B). When comparing the unstimulated cells, we observed that the human BAT selective markers, displayed a higher expression in cells isolated from the upper pole (PRDM16) and at the hilus (ZIC1) compared to subcutaneous adipocytes (Figures S4B). This expression pattern indicated that the adipogenic progenitors isolated from the perirenal depot retained a BAT phenotype in vitro. Intriguingly, LHX8, another BAT marker expressed in supraclavicular tissue and cells, was lower expressed in the perirenal cells than in the subcutaneous cells (Figure S4C), suggesting a molecular difference between supraclavicular and perirenal BAT. This observation furthermore emphasizes the need of functional validation of markers for human BAT. None of the other investigated markers including CITED1, FABP4, TBX1, CIDEA, RXRγ, HOXC8 or HOXC9, were regulated in response to NE or differentially expressed between perirenal and subcutaneous adipocytes (data not shown). Thus, the perirenal-selective expression of RXRγ observed in the tissue biopsies, was not retained in vitro. To assess the BAT functionality of the human perirenal in vitro differentiated adipocytes, we utilized the Seahorse Bioscience XF96 platform and measured the NE-induced uncoupled respiration at day 12 of differentiation. Measurements of basal and NE-induced oxygen consumption rates (OCR) were performed in the same subset of cell cultures that were characterized by flow cytometry (i.e. four cell cultures from each anatomical region, 20 cell cultures in total). Examples of OCR tracings from a subcutaneous cell culture (Figure 4C left panel) and a perirenal cell culture (Figure 4C right panel) are presented. Basal OCR was measured for 30 minutes and did not differ between regions. Following NE or saline injection, OCR was recorded for 60 minutes and a NE-dependent induction of OCR was observed in adipocytes from all perirenal regions, but not in subcutaneously derived adipocytes (Figure 4D left panel). To adjust for the slight decrease in OCR during the first 90 minutes, comparisons were made between the NE treated and saline treated wells at 60 minutes. Following addition of oligomycin (an ATP synthase inhibitor), the NE-induced increase in OCR remained significant in the perirenal adipocytes, clearly suggesting that this increase in OCR was due to uncoupled respiration. (Figure 4D right panel). Taken together, we conclude that brown fat progenitors, which can differentiate into functional brown adipocytes, are present in all assessed regions of the perirenal fat.

**Figure 4.**
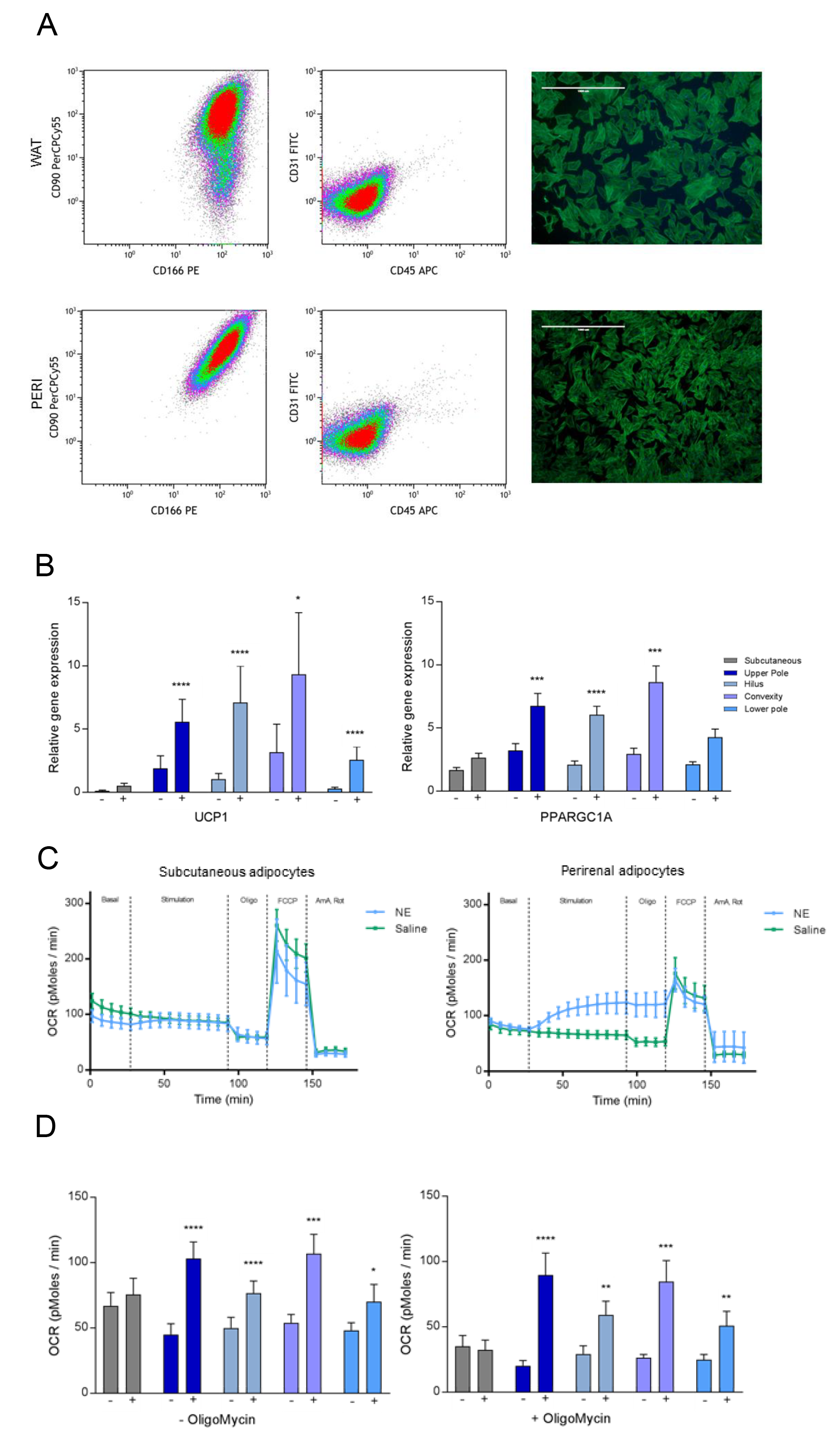
A brown fat phenotype of fat progenitors derived from perirenal fat of adults. **(A)** The first two columns show representative density plots from FACS analysis including cells labeled with CD90, CD166, CD31 and CD45. In the first column are the adipogenic cell surface markers CD90 and CD166 plotted for subcutaneous and perirenal pre-adipocytes in each row. In the second column are CD31 (endothelial cell surface marker) and CD45 (hematopoietic cell surface marker) plotted to visualize a clean cell population. See Figure S4 for gating strategy. Images (4X) in the third columns shows pre-adipocytes stained with fluorescently-labeled phalloidin and DAPI to visualize cell shape, size and confluence immediately before the FACS analysis (scale bar = 1000 μm). **(B)** Genes induced with NE in adipocytes from perirenal, but not from subcutaneous regions. Data are presented as means with SEM, comparison of expression levels was calculated using a mixed model ANOVA with Bonferroni correction and a P value below 0.05 was considered significant. * = P < 0.05, *** = P < 0.001 ****=0.0001. Subq N = 13, upper pole N = 23, hilus N =20, convexity N = 13 and lower pole N = 17. Also see figure S4B and S4C. **(C)** Example of seahorse experiments with representative tracing of a subcutaneous cell ID (Left panel) and a perirenal cell ID (right panel), showing measurements of basal respiration sequentially followed by addition of saline / 1 μM of NE stimulation, of the ATP synthase inhibitor oligomycin, the ionophore FCCP and the electron transport chain inhibitors antimycin A and rotenone. **(D)** Comparison of basal and NE induced respiration in subcutaneous and perirenal adipocytes before (left) and after (right) addition of the ATP synthase blocker oligomycin. N= 4 cell IDs pr. region. Data are presented as means with SEM. * = significant increase in oxygen consumption rate (OCR) in response to NE compared to saline treated cells within the same region. Comparisons were calculated using a mixed model ANOVA and a p-value below 0.05 was considered significant. * = p < 0.05, ** = P < 0.01, **** = p < 0.0001.

## Discussion

We here propose that the development of a dormant BAT state in the perirenal adipose is related to a decreased sympathetic activity. We addressed this by comparing adipose samples from different regions around the kidney, while taking advantage of the asymmetric sympathetic input and local sympathetic activity. Our data suggested a higher local sympathetic activity close to the adrenal gland and the hilus, which coincided with an accumulation of multilocular BAT. Transcriptomic analysis suggested reduced mitochondrial respiration in the unilocular samples, while brown fat progenitors were present regardless of location. SPARC increased in dormant BAT whereas CLSTN3 was identified as a marker for multilocular BAT.

Although PET/CT-scans combined with cooling were not applied in the current study, previous PET/CT studies have reported the most pronounced signal of active perirenal BAT to occur closest to the upper pole of the kidney^56,57^. A possible explanation to this observation could be that sympathetic nerves highly innervate, and supply NE to the endothelium of blood vessels, and both the adrenal gland and the area near the hilus of the kidney are highly vascularized. This idea is supported by anatomical studies investigating the distribution of sympathetic fibers in relationship to renal vascularization, demonstrating that 90.5 % of all nerves were found within 2.0 mm from the lumen of the renal arteries^58^.

It is clear from studies performing PET/CT-scan based screenings of subjects with different age spans, that the amount of functionally competent BAT is less with increasing age^13, 59–62^. Our subjects were healthy kidney donors, mainly around 50-60 years of age which, according to previous reports^13,59^ would be expected to have low, if any, prevalence of detectable active BAT. Interestingly, a recent study provided an FDG-PET/CT scan-based atlas, supporting the possibility of high amounts of inactive or “dormant” BAT in anatomically defined fat depots where BAT was abundant during childhood^16,63^.

Taken together with our current observations that unilocular BAT is UCP1 positive and contains brown fat precursor cells, these findings give reason to reconsider the suggestion that human BAT is beige^20,22^ or that a trans-differentiation from brown to white fat cells occur^32^. Our data rather indicate a transformation of an active BAT state to a dormant BAT state, maintaining UCP1 protein expression, a transcriptome distinct from subcutaneous WAT, and an intact pool of brown fat precursor cells. Thus, we propose that the heterogeneity often observed in human brown fat might be due to sympathetic activity-dependent shifts between active and dormant BAT states.

An important question is whether dormant BAT can be reactivated. An example where browning occurs in humans has been observed during Pheochromocytoma, a norepinephrine-producing tumor on the adrenal gland ^32,35^. Here, a ‘selective’ browning occurs, enlarging the areas were active BAT has been observed previously, identifying these regions as major targets for in vivo browning in humans. In a case study, a PET/CT-scanned individual with Pheochromocytoma was determined to possess activated BAT in the entire perirenal fat depot as well as in the nearby surrounding fat^35^. Likewise, the BAT in the supraclavicular and intercostal regions was substantially increased^35^.

Establishing how this dormant BAT is regulated will provide basis for increased understanding of how to re-activate this tissue, an attractive achievement in obesity research. We identified a secreted protein; SPARC, which was up-regulated in the dormant BAT compared to the multilocular, potentially active, BAT. It is increasingly acknowledged that BAT secretes factors, i.e. batokines with local or central effects^64,65^. Previous literature have proposed that SPARC regulate the accumulation of adipose tissue^49^, and demonstrate an opposite expression pattern to UCP1 in BAT during hibernation^53^. In the light of our data, it is possible that SPARC is involved in maintaining a dormant BAT phenotype during the absence of sympathetic activation, while the other candidate gene we identified, CLSTN3 rather associated with an active BAT state.

In conclusion, we here demonstrate that most of the perirenal fat in adult humans consist of dormant BAT with small amounts of active BAT morphology and gene expression signature close to local sources of sympathetic activity. Our data suggest that BAT in adult humans is more abundant than previously anticipated and our comparative transcriptomic analysis provides a resource for identification of novel regulators of BAT activation. Targeting such regulators might prove an efficient strategy to re-activate BAT in adult humans and restore a healthy metabolic regulation.

## Experimental Procedures

### Subjects

20 Subjects (11 men and 9 women) were included prior to scheduled kidney donation through the outpatient clinic of the Nephrological Department, Rigshospitalet after providing informed written consent. Participants were included throughout the period of June 2014 and March 2015. All subjects were examined in accordance with the custom kidney donor screening programme and were thus healthy. Anthropometrics and clinical measurements were obtained from the patient files or during the patient interview prior to participation. No additional examinations were performed and there were no additional exclusion criterions if subjects were considered eligible for donation. The Scientific-Ethics Committee of the Capital region of Copenhagen approved the study protocol, journal number H-1-2013-144 and the study was performed in accordance with the Helsinki declaration.

### Biopsies

Adipose tissue biopsies were collected during nephrectomy. Samples were collected from 3-7 areas of the perirenal fat layer surrounding the kidney. The number and specific location of the biopsies depended on the explicit fat distribution in each subject. Furthermore, a subcutaneous fat biopsy was collected from the incision site at the abdomen of all subjects. Samples were divided into 3 parts, the part for mRNA analyses was immediately snap frozen in liquid nitrogen and stored at −80 degrees until analyses were performed, while samples for immunohistochemistry were placed in formalin until further processing. Perirenal fat samples were obtained from both inside and outside Gerota’s fascia (Figure 1A) at the level of the 2 kidney poles and the convexity, whereas the hilus as such is located inside the fascia. However, since no systematic differences between samples from inside and outside the fascia were observed, samples were pooled into the 4 anatomical regions described in the result section. Three tissue samples were excluded from the analyses due to unspecific registration of their localization. Six supraclavicular samples from a previously published article ^22^ were included as reference material in the analyses of adrenergic receptor expression.

### Transcriptional profiling of human multilocular and unilocular BAT and subcutaneous WAT

A subset of the surgical biopsies described above was selected for RNA sequencing. Three groups (two perirenal and one subcutaneous) of n=4 samples were selected. The samples for the two perirenal groups had either multilocular morphology or unilocular morphology and were both positive for UCP1 protein. To increase the likelihood that the unilocular sample for RNA analysis did not contain multilocular adipocytes, we selected unilocular samples with low UCP1 expression. The subcutaneous samples were from the same individual as the unilocular samples. RNA was isolated using Trizol according to the manufacturer’s recommendations and quality control was performed using Nanodrop and Bioanalyzer. RNA sequencing (PE 100) was performed on a HiSeq 4000 platform after Truseq cDNA library construction on poly-A tail filtered RNA (BGI, Copenhagen, Denmark). The transcriptome data from 12 tissues samples were processed and analyzed by RNA-seq analysis pipeline from CLC Genomics workbench 11 (https://www.qiagenbioinformatics.com/). The human genome reference hg38 were used for the mapping and annotation with the default setting. We choose the *total counts* options for calculating the expression values. For the differential expression analysis, we used all groups pairs comparison including the FDR < 0.01 as cut off. The Gene Ontology enrichment analysis on the differentially expressed (DE) genes was performed using the clusterProfiler R package^66^. We used 0.05 q-value cut off for the term enrichment of the DE genes.

### Mouse model for dormant BAT

All experiments were approved by the Animal Ethics Committee of the North Stockholm region. Metabolic data including e.g. body weight, body composition, food intake, glucose tolerance and insulin tolerance from the mouse study have been published previously^52^. Briefly, 12-week-old male mice were single-caged at thermoneutrality (30 °C) in a 12:12-h light–dark cycle regime for at least 25 weeks. Mice were randomized in two groups given either chow (R70, Lactamin) (n=5) or high fat diet (45% calories from fat, Research Diets D12451) (n=5), ad libitum. Mice were sacrificed by CO2 anaesthesia. Interscapular BAT and inguinal white adipose tissue was dissected, snap-frozen in liquid nitrogen and stored in −80°C until further analysis. Total RNA was extracted using TRI reagent and cDNA synthesis and quantitative real-time PCR was performed as previously described^52^. The primer sequences are presented in supplementary material.

### Isolation, culture and differentiation of human adipogenic progenitor cells

Primary cell cultures were established as previously described ^22^ Adipogenic progenitor cells were isolated from the stromal vascular fraction of the biopsies on the day they were obtained. Biopsies were collected in DMEM/F12 (Gibco) with 1% penicillin / streptomycin (PS; life technologies) and tubes were kept on ice during transport from the operating room to the cell-lab. Biopsies were digested in a buffer containing 10 mg collagenase II (C6885-1G, Sigma) and 100 mg BSA (A8806- 5G, Sigma) in 10 ml DMEM/F12 for 20-30 minutes at 37° C while gently shaken. Following digestion, the suspension was filtered through a cell strainer (70 μm size) and cells were left to settle for 5 minutes before the layer below the floating, mature adipocytes was filtered through a thin filter (30 micron). The cell suspension was centrifuged for 7 min at 800 g and the cell pellet was washed with DMEM/F12 and then centrifuged again before being resuspended in DMEM/F12, 1% PS, 10% fetal bovine serum (FBS) (Life technologies) and seeded in a 25 cm^2^ culture flask. Media was changed the day following isolation and then every second day until cells were 80% confluent, at this point cultures were split into a 10 cm dish (passage 0).

Cells were expanded by splitting 1:3. At passage 1, cells were seeded for gene expression experiments in proliferation media consisting of DMEM/F12, 10 % FBS, 1 % PS and 1 nM Fibroblast growth factor□acidic (FGF□1) (ImmunoTools). Cells were grown at 37° C in an atmosphere of 5% CO2 and the medium was changed every second day. Adipocyte differentiation was induced two days after preadipocyte cultures were 100% confluent by addition of a differentiation cocktail consisting of DMEM/F12 containing 1% PS, 0.1 μM dexamethasone (Sigma□Aldrich), 100 nM insulin (Actrapid, Novo Nordisk or Humulin, Eli Lilly), 200 nM rosiglitazone (Sigma□Aldrich), 540 μM isobutylmethylxanthine (Sigma□Aldrich), 2 nM T3 (Sigma□Aldrich) and 10 μg/ml transferrin (Sigma□Aldrich). After three days of differentiation, isobutylmethylxanthine was removed from the cell culture media, and after an additional three days rosiglitazone was removed from the media for the remaining 6 days of differentiation. On the 12^th^ day of differentiation the media was changed to DMEM/F12, 1% PS for 2 hours before stimulation with 10 μM norepinephrine (Sigma Aldrich, A9512 L-(−)-Norepinephrine (+)-bitartrate salt monohydrate diluted in sterile H2O) or sterile H2O. After 4 hours of stimulation, cells were harvested using TRizol (Invitrogen) and stored on −80° C until PCR analyses were performed.

The degree of cell differentiation was evaluated based on a combination of subjective evaluation of the amount of accumulated lipid droplets (% of culture) on the day of stimulation and FABP4 mRNA expression (Figure S3). For seven of the cultures no visible estimation had been registered and thus only FABP4 expression was used for the estimation in these cultures.

On the final day of differentiation cultures were stimulated with either 10 μM of NE or vehicle treatment, before being harvested for gene expression analyses. Primary cell cultures were established from all participants, but only cultures from 18 of the subjects were included in the study, total cell ID N = 86, (13 subcutaneous and 73 perirenal). Cultures from 2 subjects were excluded due to infection and one culture was excluded based on complete lack of differentiation. The availability of the cell cultures for other research groups is dependent on specific permission from the Danish Data Protection Agency and on the researcher’s adherence to the specifications of this permission.

### RNA Isolation and Quantitative Real time PCR

Total RNA isolation from adipose tissue biopsies was performed using TRizol reagent according to the manufacture’s protocol. RNA was dissolved in nuclease-free water and quantified using a Nanodrop ND 1000 (Saveen Biotech). Total RNA (0.25μg) was reverse-transcribed using the High Capacity cDNA Reverse Transcription Kit (Applied Biosystems). cDNA samples were loaded in triplicate and qPCR was performed using Real Time quantitative PCR, using the ViiA^tm^ 7 platform (Applied Biosystems). Relative quantification was conducted by either SYBRgreen fluorescent dye (Applied Biosystems) or TaqMan Gene Expression Assays (Applied Biosystems). All procedures were performed according to the manufacturer’s protocol. Target mRNA expression was calculated based on the standard curve method and was normalised to the reference gene PPIA. For primer sequences see table S2 and S3.

### Immunohistochemistry

Immunohistochemical analyses were performed at the Pathological Department, Rigshospitalet. The tissue samples were placed in 10% neutral-buffered formalin and processed routinely. After fixation for about 24 h, the tissue was placed in cassettes and transferred to a Leica Peloris Rapid Tissue Processor (Leica Biosystems GmbH, Nussloch, Germany). The tissue was then paraffin-embedded and sectioned manually, and stained with haematoxylin-eosin (HE). The immunohistochemical reactions were performed using a BenchMark ULTRA IHC/ISH Staining Module with Optiview visualization reagents (Ventana Medical Systems Inc., Tucson, Arizona, USA). Immunostainings were performed using: UCP1: Rabbit-polyclonal UCP1-antibody (Abcam), dilution 1:500, pretreatment ph9 (32 minutes) + protease3 (4 minutes), RXRγ: Rabbit-polyclonal RXRγ-antibody (Abcam), dilution 1:2000, pretreatment pH9 (32 minutes), PRDM16: Rabbit-polyclonal antibody (Abcam), diluation, 1:500, pretreatment pH6 (32 minutes). Sections were evaluated as follows: multilocular appearance in the HE staining and UCP1 positive (MBAT), HE unilocular, but UCP1 positive (UBAT) or both HE unilocular and UCP1 negative (WAT). PRDM16 (nuclear) were scored as either positive or negative, i.e. if there were any positive (nuclear reaction) in any of the cells present. No attempt was made to quantitate these results. RXRγ stainings (nuclear) were scored as positive or with only few positive cells. For UCP1 stainings a hibernoma tissue section was included as positive and negative control. Samples were evaluated by the author SD.

### Protein expression

Perirenal and subcutaneous protein expression was assessed by Western blotting. Primary antibodies were used at the following concentrations: UCP1 (Abcam ab155117; 1:1000), tyrosine hydroxylase (Abcam ab112; 1:200) and OXPHOS (Abcam ab110413; 1:1000). Primary antibodies were detected with either anti-rabbit or anti-mouse horseradish peroxidase-linked IgG (Dako) at a concentration of 1:5000 and imaged using Supersignal West Femto (Pierce). Data are expressed relative to total protein expression using stain free UV imaging (Biorad) and normalized to a pooled perirenal sample included on all gels. Gels were quantified using Image Lab version 5.2.1 (Biorad) software.

### Flow cytometry

Flow cytometry was performed on cells from passage three. The cells were proliferated according to standard protocol in DMEM/F12 medium supplemented with 10% FBS, 1% PS, and 1 nmol/L fibroblast growth factor in 5% CO2, 37°C environment until they reached 80% confluence. Cells were harvested using TrypLE (Gibco; Life Technologies) followed by a wash in buffer (PBS containing 2% FBS and 0.01% NaN3) and were afterwards resuspended in staining buffer (PBS containing 2% FBS, 1% Human Serum [catalog #1001291552; Sigma] and 0.01% NaN3). Anti-human CD45-APC, CD31-FITC, CD90- PerCP.Cy5.5, and CD166-PE (BD Pharmingen), antibodies were added to the cells and flow cytometry was applied for quantification using a FACS Fortessa (BD Bioscience). For compensation, single stain was used with one drop of negative control beads and anti-mouse IgG beads (BD Biosciences). Data analysis was performed using Kaluza software, version 1.2 (Beckman Coulter).

### Cell Imaging

When pre-adipocytes reached a confluence at 80% they were fixated in 4% formaldehyde (15 min) and then permeabilized in 0.5% Triton X-100 (15 min). F-actin was then labelled with ActinGreen™ 488 ReadyProbes^®^ (Thermo Fisher) for 30 min and DAPI (Thermo Fisher) for visualization of the nucleus. Imaging was performed with EVOS FL Cell Imaging System (Thermo Fisher.

### Oxygen consumption analyses

Oxygen consumption was measured using a Seahorse Bioscience XF96 Extracellular Flux Analyser according to the manufacturer’s protocol. Adipocytes were grown until reaching 100% confluency and were then seeded in seahorse plates at a 1:1 ratio and differentiated as described above. Experiments were performed on day 12 of differentiation on cells in passage three. Oxygen consumption rate was assessed in 20 primary adipocyte cultures, four per region. The sample set was selected based on approximately equal degree of differentiation. The results were extracted from the Seahorse Program Wave 2.2.0. Baseline measurements of OCR were performed for 30 minutes before NE or saline was added and measurements of the concomitant responses were recorded for 60 minutes. All other states were induced using the seahorse XF cell mito stress test kit according to the manufactures protocol. After 90 minutes, leak state was induced by adding Oligomycin, which inhibits the ATP synthase. Leak state measurements were performed for 20 minutes, then the ionophore (carbonyl cyanide-4-(trifluoromethoxy) phenylhydrazone) (FCCP), which collapses the proton gradient across the mitochondrial inner membrane resulting in a completely uncoupled state. After an additional 20 minutes Antimycin A and Rotenone were added to inhibit complexes III and I respectively, resulting in only non-mitochondrial respiration.

For data analyses OCR was corrected for non-mitochondrial respiration as assessed by the Seahorse XF cell mitochondrial stress test kit. To adjust for a slight decrease in OCR during the first 90 minutes OCR changes were calculated as the difference between the NE treated wells and the saline treated wells and data analyses were performed. Wells were excluded from the data analyses if OCR were +/-20% of the mean in that series of replicate values. In total 17 wells were excluded based on this.

### Statistical Analyses

Statistical analyses were performed using SAS 9.4 statistical software. Data are presented as means (bars) with whiskers representing standard error of the mean (SEM), except in Figure 1A and S1 where data are presented as Tukey box plots, with boxes representing the interquartile range (IQR), whiskers representing the 25-75% percentile and data points > 1,5 times above IQR is presented as individual data points in order to present the large variation adequately, and in Fig. 2C., where data are presented as means with whiskers representing minimum (min) – maximum (max) values. For brown, brite / beige and white fat markers, differences in gene expression were calculated using repeated measures mixed models ANOVA with Tukey or Bonferroni correction and a P-value below 0.05 was considered significant (Littell, et al 2006). For receptor analyses, differences in gene expression were calculated using mixed model ANOVA comparing perirenal and supraclavicular fat to subcutaneous samples. However, in some cases mixed models analyses could not be performed, due to infinite likelihood related to a low n number of the supraclavicular samples. In these cases, (ADRA1A, ADRA1D, ADRA2A, ADRA2B and ADRA2C), mixed model analysis was used to assess the difference between subcutaneous and perirenal samples, excluding the supraclavicular samples from this analysis. A one-way ANOVA was then performed to assess differences between subcutaneous and supraclavicular samples for the mentioned receptors. In either case a P value below 0.05 was considered statistically significant. Differences in protein expression was also assessed applying a repeated measures mixed models ANOVA with the Kenward-Roger approximation (Kenward and Roger, 1997). Differences in OCR measurements were calculated using repeated measures mixed model ANOVA applying Tukey correction for between region analyses. A P-value below 0.05 was considered statistically significant. Logarithmic transformation was applied in all analyses when appropriate to attain normal distribution. Correlations between UCP1 expression and other genes, were analysed using ordinary least squares (OLS) regression analyses with manual backwards elimination of independent variables. Tables and figures were designed using GraphPad Prism 7.

## Author Contributions

CS and SN supervised the study. CS, SN, NZJ, NP, BKP, RKM: hypothesis generation, conceptual design, data analysis, and manuscript preparation. NZJ, AF, ESA, SH, HBH, SN, NP, SD, PB, BFR, HSS, NSH: conducting experiments and data analysis. All authors edited and approved the final manuscript.

## Acknowledgements

The authors sincerely thank the study participants. We furthermore thank the staff at the Departments of Nephrology and Urology, Rigshospitalet for their invaluable assistance with inclusion of participants and collection of data with specific acknowledgement for the efforts of senior consultant Søren Schwartz Sørensen and Ph.d-student Lene Kjær Olsen. Noemi G. James, Maria M. Scheel and Lone Peijs are acknowledged for technical assistance. Pernille Frederiksen at the Department of Pathology is acknowledged for technical assistance with the immunohistochemistry. We are also thankful to the Department of Biostatistics at University of Copenhagen for their statistical advisory service.

The Centre for Physical Activity Research (CFAS) is supported by a grant from TrygFonden. During the study period the Centre for Inflammation and Metabolism (CIM) was supported by a grant from the Danish National Research Foundation (DNRF55). The Novo Nordisk Foundation Center for Basic Metabolic Research (http://www.metabol.ku.dk) is supported by an unconditional grant from the Novo Nordisk Foundation to University of Copenhagen. The study was further supported by research grants from the Lundbeck foundation (CS), the Danish Diabetes Academy supported by the Novo Nordisk Foundation, The University of Copenhagen, the Research foundation of Rigshospitalet, Carl and Ellen Hertz Foundation, The Foundation of 1870, Oda and Hans Svenningsens Foundation, The Foundation of the Family Hede Nielsen. CIM/CFAS is a member of DD2 - the Danish Center for Strategic Research in Type 2 Diabetes (the Danish Council for Strategic Research, grant no. 09-067009 and 09-075724).

## Disclosure of limited availability of biological material

We hereby disclose that the availability of biological material including the human cell cultures is dependent on specific permission from the Danish Data Protection Agency and on the researcher’s adherence to the specifications of this permission.

## References

1. Cypess, A. M. et al. Identification and Importance of Brown Adipose Tissue in Adult Humans. N. Engl. J. Med. 360, 1509–1517 (2009).

2. Nedergaard, J., Bengtsson, T. & Cannon, B. Unexpected evidence for active brown adipose tissue in adult humans. Am. J. Physiol. Endocrinol. Metab. 293, 444–452 (2007).

3. Saito, M. et al. High Incidence of Metabolically Active Brown Adipose Effects of Cold Exposure and Adiposity. Diabetes 58, 1526–1531 (2009).

4. Van Marken Lichtenbelt, W. D. et al. Cold-activated brown adipose tissue in healthy men. N. Engl. J. Med. 360, 1500–1508 (2009).

5. Virtanen, K. A. et al. Functional Brown Adipose Tissue in Healthy Adults. N. Engl. J. Med. 360, 1518–25 (2009).

6. Zingaretti, M. C. et al. The presence of UCP1 demonstrates that metabolically active adipose tissue in the neck of adult humans truly represents brown adipose tissue. FASEB J. 23, 3113–3120 (2009).

7. Cannon, B. & Nedergaard, J. Brown Adipose Tissue: Function and Physiological Significance. Physiol. Rev. 84, 277–359 (2004).

8. Cypess, A. M. et al. Activation of Human Brown Adipose Tissue by a β3-Adrenergic Receptor Agonist. Cell Metab. 21, 33–38 (2015).

9. Hanssen, M. J. W. et al. Short-term cold acclimation improves insulin sensitivity in patients with type 2 diabetes mellitus. Nat. Med. 21, 6–10 (2015).

10. Chondronikola, M. et al. Brown Adipose Tissue Improves Whole-Body Glucose Homeostasis and Insulin Sensitivity in Humans. Diabetes 63, 4089–4099 (2014).

11. Lee, P. et al. Temperature-Acclimated Brown Adipose Tissue Modulates Insulin Sensitivity in Humans. Diabetes 63, 3686–3698 (2014).

12. Hanssen, M. J. W. et al. Short-term cold acclimation recruits brown adipose tissue in obese humans. Diabetes (2016). doi:10.2337/db15-1372

13. Yoneshiro, T. et al. Age-related decrease in cold-activated brown adipose tissue and accumulation of body fat in healthy humans. Obesity (Silver Spring). 19, 1755–60 (2011).

14. Yoneshiro, T. et al. Impact of UCP1 and β3AR gene polymorphisms on age-related changes in brown adipose tissue and adiposity in humans. Int. J. Obes. (Lond). 37, 993–998 (2013).

15. Matsushita, M. et al. Impact of brown adipose tissue on body fatness and glucose metabolism in healthy humans. Int. J. Obes. 38, 812–817 (2014).

16. Heaton, J. M. The distribution of brown adipose tissue in the human. 35–39 (1972).

17. Tanuma, Y., Tamamoto, M., Ito, T. & Yokochi, C. The occurrence of brown adipose tissue in perirenal fat in Japanese. Arch. Histol. Jpn. 38, 43–70 (1975).

18. Seale, P. et al. Prdm16 determines the thermogenic program of subcutaneous white adipose tissue in mice. J. Clin. Invest. 121, 96–105 (2011).

19. Petrovic, N. et al. Chronic peroxisome proliferator-activated receptor gamma (PPARG) activation of epididymally derived white adipocyte cultures reveals a population of thermogenically competent, UCP1-containing adipocytes molecularly distinct from classic brown adipocytes. J. Biol. Chem. 285, 7153–7164 (2010).

20. Wu, J. et al. Beige Adipocytes Are a Distinct Type of Thermogenic Fat Cell in Mouse and Human. Cell 150, 366–376 (2012).

21. Lidell, M. E. et al. Evidence for two types of brown adipose tissue in humans. Nat. Med. 19, 631–634 (2013).

22. Jespersen, N. Z. et al. A classical brown adipose tissue mrna signature partly overlaps with brite in the supraclavicular region of adult humans. Cell Metab. 17, (2013).

23. Xue, R. et al. Clonal analyses and gene profiling identify genetic biomarkers of the thermogenic potential of human brown and white preadipocytes. Nat. Med. 21, 760–768 (2015).

24. Li, X. et al. Determination of UCP1 expression in subcutaneous and perirenal adipose tissues of patients with hypertension. Endocrine (2015). doi:10.1007/s12020-015-0572-3

25. Van Den Beukel, J. C. et al. Women have more potential to induce browning of perirenal adipose tissue than men. Obesity 23, n/a-n/a (2015).

26. Nagano, G. et al. Activation of Classical Brown Adipocytes in the Adult Human Perirenal Depot Is Highly Correlated with PRDM16–EHMT1 Complex Expression. PLoS One 10, e0122584 (2015).

27. Betz, M. J. et al. Presence of brown adipocytes in retroperitoneal fat from patients with benign adrenal tumors: Relationship with outdoor temperature. J. Clin. Endocrinol. Metab. (2013). doi:10.1210/jc.2012-3535

28. Svensson, P. A. et al. Characterization of brown adipose tissue in the human perirenal depot. Obesity 22, 1830–1837 (2014).

29. Vergnes, L. et al. Adipocyte Browning and Higher Mitochondrial Function in Periadrenal But Not SC Fat in Pheochromocytoma. J Clin Endocrinol Metab 101, 4440–4448 (2016).

30. Goldstein, D. S., Eisenhofer, G. & Kopin, I. J. Sources and significance of plasma levels of catechols and their metabolites in humans. J. Pharmacol. Exp. Ther. 305, 800–11 (2003).

31. Dundamadappa, S. K. et al. Imaging of brown fat associated with adrenal pheochromocytoma. Acta Radiol. 48, 468–72 (2007).

32. Frontini, A. et al. White-to-brown transdifferentiation of omental adipocytes in patients affected by pheochromocytoma. Biochim. Biophys. Acta 1831, 950–959 (2013).

33. Puar, T. et al. Genotype-dependent brown adipose tissue activation in patients with pheochromocytoma and paraganglioma. J. Clin. Endocrinol. Metab. (2016). doi:10.1210/jc.2015-3205

34. Rona, G. Changes in Adipose Tissue Accompanying Pheochromocytoma. Can. Med. Assoc. J. 91, 303–5 (1964).

35. Søndergaard, E. et al. Chronic adrenergic stimulation induces brown adipose tissue differentiation in visceral adipose tissue. Diabet. Med. 32, e4–e8 (2015).

36. de Jong, J. et al. The β3-adrenergic receptor is dispensable for browning of adipose tissues. Am J Physiol Endocrinol Metab. 1;312, E508–E518

37. Susulic, V. S. et al. Targeted disruption of the beta 3-adrenergic receptor gene. J. Biol. Chem. 270, 29483–92 (1995).

38. Sharp, L. Z. et al. Human BAT possesses molecular signatures that resemble beige/brite cells. PLoS One 7, e49452 (2012).

39. Puigserver, P. et al. A cold-inducible coactivator of nuclear receptors linked to adaptive thermogenesis. Cell 92, 829–839 (1998).

40. de Jong, J. M. A., Larsson, O., Cannon, B. & Nedergaard, J. A stringent validation of mouse adipose tissue identity markers. Am. J. Physiol. - Endocrinol. Metab. 308, E1085–E1105 (2015).

41. Timmons, J. a et al. Myogenic gene expression signature establishes that brown and white adipocytes originate from distinct cell lineages. Proc. Natl. Acad. Sci. U. S. A. 104, 4401–4406 (2007).

42. Seale, P. et al. PRDM16 controls a brown fat/skeletal muscle switch. Nature 454, 961–967 (2008).

43. Kajimura, S. et al. Initiation of myoblast to brown fat switch by a PRDM16–C/EBP-β transcriptional complex. Nature 460, 1154–1158 (2009).

44. Seale, P. et al. Transcriptional Control of Brown Fat Determination by PRDM16. Cell Metab. 6, 38–54 (2007).

45. Mori, M., Nakagami, H., Rodriguez-Araujo, G., Nimura, K. & Kaneda, Y. Essential role for miR-196a in brown adipogenesis of white fat progenitor cells. PLoS Biol. 10, e1001314 (2012).

46. Kazak, L. et al. A Creatine-Driven Substrate Cycle Enhances Energy Expenditure and Thermogenesis in Beige Fat Article A Creatine-Driven Substrate Cycle Enhances Energy Expenditure and Thermogenesis in Beige Fat. Cell 163, 643–655 (2015).

47. Müller, S. et al. Proteomic Analysis of Human Brown Adipose Tissue Reveals Utilization of Coupled and Uncoupled Energy Expenditure Pathways. Sci. Rep. 6, 30030 (2016).

48. Shinoda, K. et al. Genetic and functional characterization of clonally derived adult human brown adipocytes. Nat. Med. 21, 389–394 (2015).

49. Bradshaw, A. D., Graves, D. C., Motamed, K. & Sage, E. H. SPARC-null mice exhibit increased adiposity without significant differences in overall body weight. Proc. Natl. Acad. Sci. U. S. A. 100, 6045–50 (2003).

50. Fischer, A. W. et al. UCP1 inhibition in Cidea-overexpressing mice is physiologically counteracted by brown adipose tissue hyperrecruitment. Am. J. Physiol. - Endocrinol. Metab. 312, E72–E87 (2017).

51. Sanchez-Gurmaches, J. et al. Brown Fat AKT2 Is a Cold-Induced Kinase that Stimulates ChREBP-Mediated De Novo Lipogenesis to Optimize Fuel Storage and Thermogenesis. Cell Metab. 27, 195–209 (2018).

52. Abreu-Vieira, G. et al. Cidea improves the metabolic profile through expansion of adipose tissue. Nat. Commun. 6, (2015).

53. Yan, J. et al. Detection of differential gene expression in brown adipose tissue of hibernating arctic ground squirrels with mouse microarrays. 346–353 (2006). doi:10.1152/physiolgenomics.00260.2005.

54. Peng, B., Gu, Y., Xiong, Y., Zheng, G. & He, Z. Microarray-Assisted Pathway Analysis Identifies MT1X & NF??B as Mediators of TCRP1-Associated Resistance to Cisplatin in Oral Squamous Cell Carcinoma. PLoS One 7, (2012).

55. Schultz, N. et al. Impaired leptin gene expression and release in cultured preadipocytes isolated from individuals born with low birth weight. Diabetes 63, 111–121 (2014).

56. Bar-Shalom, R., Gaitini, D., Keidar, Z. & Israel, O. Non-malignant FDG uptake in infradiaphragmatic adipose tissue: a new site of physiological tracer biodistribution characterised by PET/CT. Eur. J. Nucl. Med. Mol. Imaging 31, 1105–1113 (2004).

57. Yeung, H. W. D., Grewal, R. K., Gonen, M., Schöder, H. & Larson, S. M. Patterns of (18)F-FDG uptake in adipose tissue and muscle: a potential source of false-positives for PET. J. Nucl. Med. 44, 1789–96 (2003).

58. Atherton, D.., Deep, N.. & Mendelsohn, F. . Micro-anatomy of the renal sympathetic nervous system: a human postmortem histologic study. Clin. Anat. 25, 628–33 (2012).

59. Cronin, C. G. et al. Brown Fat at PET/CT: Correlation with Patient Characteristics. Radiology 263, 836–842 (2012).

60. Pfannenberg, C. et al. Impact of age on the relationships of brown adipose tissue with sex and adiposity in humans. Diabetes 59, 1789–1793 (2010).

61. Ouellet, V. et al. Outdoor temperature, age, sex, body mass index, and diabetic status determine the prevalence, mass, and glucose-uptake activity of 18F-FDG-detected BAT in humans. J. Clin. Endocrinol. Metab. 96, 192–199 (2011).

62. Gerngroß, C., Schretter, J., Klingenspor, M. & Schwaiger, M. Active brown fat during 18 FDG-PET / CT imaging defines a patient group with characteristic traits and an increased probability of brown fat redetection. J. Nucl. Med. 1–30 (2017).

63. Leitner, B. P. et al. Mapping of human brown adipose tissue in lean and obese young men. Proc. Natl. Acad. Sci. U. S. A. 114, 6–11 (2017).

64. Villarroya, F., Cereijo, R., Villarroya, J. & Giralt, M. Brown adipose tissue as a secretory organ. Nat. Rev. Endocrinol. 13, 26–35 (2016).

65. Scheele, C. & Nielsen, S. Metabolic regulation and the anti-obesity perspectives of human brown fat. Redox Biol. 12, 770–775 (2017).

66. Yu, G., Wang, L.-G., Han, Y. & He, Q.-Y. clusterProfiler: an R Package for Comparing Biological Themes Among Gene Clusters. Omi. A J. Integr. Biol. 16, 284–287 (2012).

67. Littell, Ramon C., George A. Milliken, W. W. & Stroup, Russell D. Wolfinger, and O. S. SAS for mixed models. (Cary, NC: SAS Institute Inc., 2006).

68. Kenward, M. G. & Roger, J. H. Small Sample Inference for Fixed Effects from Restricted Maximum Likelihood. Biometrics 53, 983–997 (1997).

69. Robinson, M. D. & Oshlack, A. A scaling normalization method for differential expression analysis of RNA-seq data. Genome Biol. 11, (2010).

